# D/E-rich peptides are less suitable than D/E-deficient peptides for identification by negative-ion HCD due to scarce production of sequencing ions from multiply charged precursors

**DOI:** 10.1101/2022.06.30.498346

**Authors:** Mei-Qing Zuo, Rui-Xiang Sun, Meng-Qiu Dong

## Abstract

Highly acidic, D/E-rich peptides or proteins are difficult to identify by positive-ion-mode mass spectrometry (MS), and negative-ion-mode MS is an attractive but insufficiently explored alternative. Based on high-resolution and accurate-mass MS analysis of 115 synthetic peptides of 5-28 amino acids, we confirmed that higher-energy collisional dissociation (HCD) of deprotonated peptides induced abundant backbone or side-chain neutral losses (NL), and updated the ranking list of NLs by abundance. The most abundant fragment ion types are *y-* > *x-, z-* > *c-* if the NL ions are included, or *c-* > *y-* > *z-* > *b-* if not. The most frequent side-chain NLs involve amino acids C, S, T, D, E, N, Q, and R. Although NL of CO_2_ is common for all peptides, it is markedly enhanced in D/E-containing peptides. Long peptides and D/E-rich peptides are prone to carrying multiple negative charges. HCD spectra produced from multiply deprotonated peptides have a lower fraction of sequencing ions (i.e., *a-, b-, c-, x-, y-, z-* ions) than those produced from 1^-^ ([M-H]^-^) and 2^-^ ([M-2H]^2-^) precursors. Based on the above findings, we predict that under negative-ion HCD, D/E-rich peptides should be difficult to identify and that choosing a protease to generate peptides containing fewer D/E residues will improve identification of highly acidic proteins.

## 1. Introduction

Mass spectrometry (MS)-based proteomics has matured in recent years as manifested by the deep MS analyses of the human proteome in which up to 17,100 protein groups (86.5% of the total by estimation) have been identified (*1, 2*). However, not every protein is readily identifiable, for example, highly acidic proteins tend to escape MS identification (*3*). For proteins that are amenable to MS-based proteomics analysis, the coverage of protein sequence is incomplete. In standard proteomics analysis, which is carried out in the positive ion mode, acidic peptides are harder to identify than hydrophobic peptides containing basic residues (*4*).

Theoretically, negative-ion-mode MS analysis can complement the ubiquitous positive-ion-mode analysis by improving identification of highly acidic peptides, which are difficult to ionize by positive-ion electrospray. Acidic peptides include the ones that are rich in acidic amino acid (aa) residues aspartic acid (Asp or D) or glutamic acid (Glu or E), and the ones that are heavily modified by negatively charged post-translational modifications (e.g., phosphorylation or sulfation). Because highly acidic peptides favor deprotonation, they ionize better in the negative-ion mode than in the positive-ion mode. This makes negative-ion-mode MS an attractive way for identifying those peptides. However, production of a precursor ion is only the first step; subsequent fragmentation of the precursor ion dictates identification. Although several studies have shown that MS analysis of deprotonated peptides (or interchangeably, peptide anions) yields information unattainable in the positive-ion mode (*3, 5, 6*), none addressed the question whether highly acidic peptides fragment well enough for identification in the negative-ion mode.

In fact, there are only limited studies on MS analysis of peptides in the negative-ion mode. Bowie and colleagues examined fragmentation of singly-charged peptide anions using mostly collision-induced dissociation (CID) (*7-10*) and summarized their findings in three reviews (*11-13*). Cassady and co-worker studied the fragmentation of several doubly or multiply charged peptide anions by CID (*5*). Coon and co-workers performed proteomics analysis using negative electron transfer dissociation (NETD) (*3, 6*), and the Brodbelt team did similar analysis using 193-nm ultraviolet photodissociation (UVPD) in the negative-ion mode (*14-16*). Some of these studies prove that negative-ion-mode MS can identify more acidic peptides and access part of the proteome unseen under the positive-ion mode, which includes highly acidic proteins or regions and membrane proteins (*3, 6, 15-16*). However, the numbers of peptides identified in the negative ion mode by either NETD or UVPD are a fraction of those identified in the positive ion mode. This is in part due to a lack of software tools optimized for negative-ion-mode MS data analysis, which in turn has to do with an insufficient understanding of the fragmentation behaviors of peptide anions.

Several studies have shown that fragmentation of deprotonated peptides by either CID or NETD generates an abundance of neutral loss (NL) peaks including a variety of losses from amino acid side chains (*17-22*). For example, threonine, serine, and tryptophan generate characteristic NL of 44.02621 (C_2_H_4_O), 30.01056 (CH_2_O), and 129.05785 (C_9_H_7_N) Da, respectively (*18, 19*). In a 2019 study, we show that the number of fragment ions with NL is greater than the number of fragment ions without NL, and the summed intensity of the former is much higher than the summed intensity of the latter (*23*). Therefore, peptide identification in negative-ion-mode MS analysis must take into consideration the NL peaks in MS2 spectra.

However, as the model peptides used in the previous studies are limited in one way or another, characterization of NL of deprotonated peptides is incomplete. In most studies, only a small number of synthetic peptides were analyzed, typically less than ten and occasionally several dozens. One exception is a 2017 study by Liang et al., in which 527 model peptides were analyzed, but they were very short peptides—137 had 3-6 residues and the rest were dipeptides.

In this study, we analyzed the negative-ion-mode MS2 spectra of 115 synthetic peptides of 5-28 residues. The MS2 spectra were generated using higher-energy collisional dissociation (HCD). We focused on finding the different varieties of NL peaks and the effects of acidic residues. The frequencies of neutral losses associated with different amino acids are updated, as are the occurrence frequency and relative abundance of six ion types (*a-, b-, c-, x-, y-*, and *z-*). Neutral loss of CO_2_ is one of the most prominent features seen in the HCD spectra of deprotonated peptides. D/E-containing peptides have more neural loss of CO_2_ off precursors and fragment ions than D/E-free peptides. The longer the peptide and the more D/E residues it contains, the more charges it carries in the negative-ion mode. When a peptide carries more than two negative charges, the percentage of sequencing ions in HCD spectra decreases. This suggests that for peptide sequencing by negative-ion HCD, highly acidic, D/E-rich peptides are less suitable than D/E-deficient peptides. Therefore, for highly acidic proteins or regions of proteins, sample digestion using a protease that cuts at D or E will be a better choice than trypsin, which cuts at K or R, as the resulting tryptic peptides will tend to be long and contain multiple D/E residues.

## 2. Materials and methods

### 2.1 Materials and sample preparation

A toal of 12 sequence-matched peptides (Table 1) were synthesized in three groups by Guoping Chemical Ltd. (Hefei, China). The concentrations of the peptide stock solutions were measured as described (*24*) using NanoDrop One (Thermo scientific) at 205 nm and the extinction coefficients listed in Table 1. For the MS1 analysis experiment in Figure 1, the four peptides in the same group were combined together to similar concentrations. For the group 1 and group 3 peptides (AAAAAD/E/K/R, GYWGTNLGQPHSLATD/E/K/R), each was diluted with 7.5 mM hexylamine (Energy chemical, Shanghai, China), 50% acetonitrile (Thermo Scientific) and sprayed into the mass spectrometer using a syringe pump at the concentrations listed in Table 1. Because the four cysteine-containing peptides of group 2 (IAVNYCMFGD/E/K/R) formed disulfide-bonded dimers, they were reduced using 5 mM TCEP (tris(2-carboxyethyl)phosphine), alkylated using 10 mM iodoacetamide, and then sprayed into the mass spectrometer as a mixture. For Figure 2 and Figure 3, the MS2 spectra of individual peptides were all acquired from a sample containing only one pepttide.

**Table 1.**
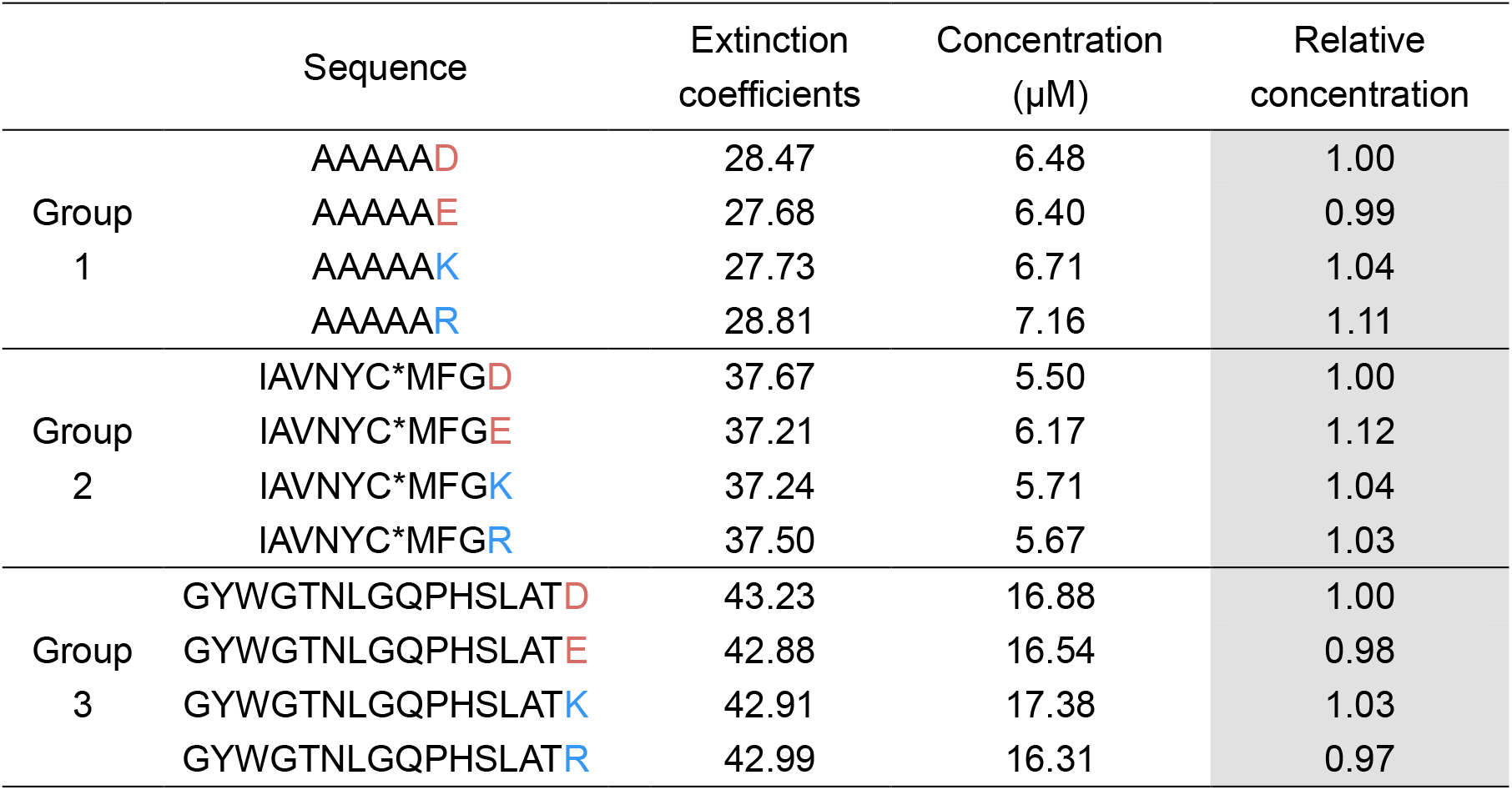
Twelve synthetic peptides used to compare the effects of C-terminal residues

|  | Sequence | Extinction coefficients | Concentration (μM) | Relative concentration |
| --- | --- | --- | --- | --- |
| Group 1 | AAAAAD | 28.47 | 6.48 | 1.00 |
|  | AAAAAE | 27.68 | 6.40 | 0.99 |
|  | AAAAAK | 27.73 | 6.71 | 1.04 |
|  | AAAAAR | 28.81 | 7.16 | 1.11 |
| Group 2 | IAVNYC*MFGD | 37.67 | 5.50 | 1.00 |
|  | IAVNYC*MFGE | 37.21 | 6.17 | 1.12 |
|  | IAVNYC*MFGK | 37.24 | 5.71 | 1.04 |
|  | IAVNYC*MFGR | 37.50 | 5.67 | 1.03 |
| Group 3 | GYWGTNLGQPHSLATD | 43.23 | 16.88 | 1.00 |
|  | GYWGTNLGQPHSLATE | 42.88 | 16.54 | 0.98 |
|  | GYWGTNLGQPHSLATK | 42.91 | 17.38 | 1.03 |
|  | GYWGTNLGQPHSLATR | 42.99 | 16.31 | 0.97 |

**Figure 1.**
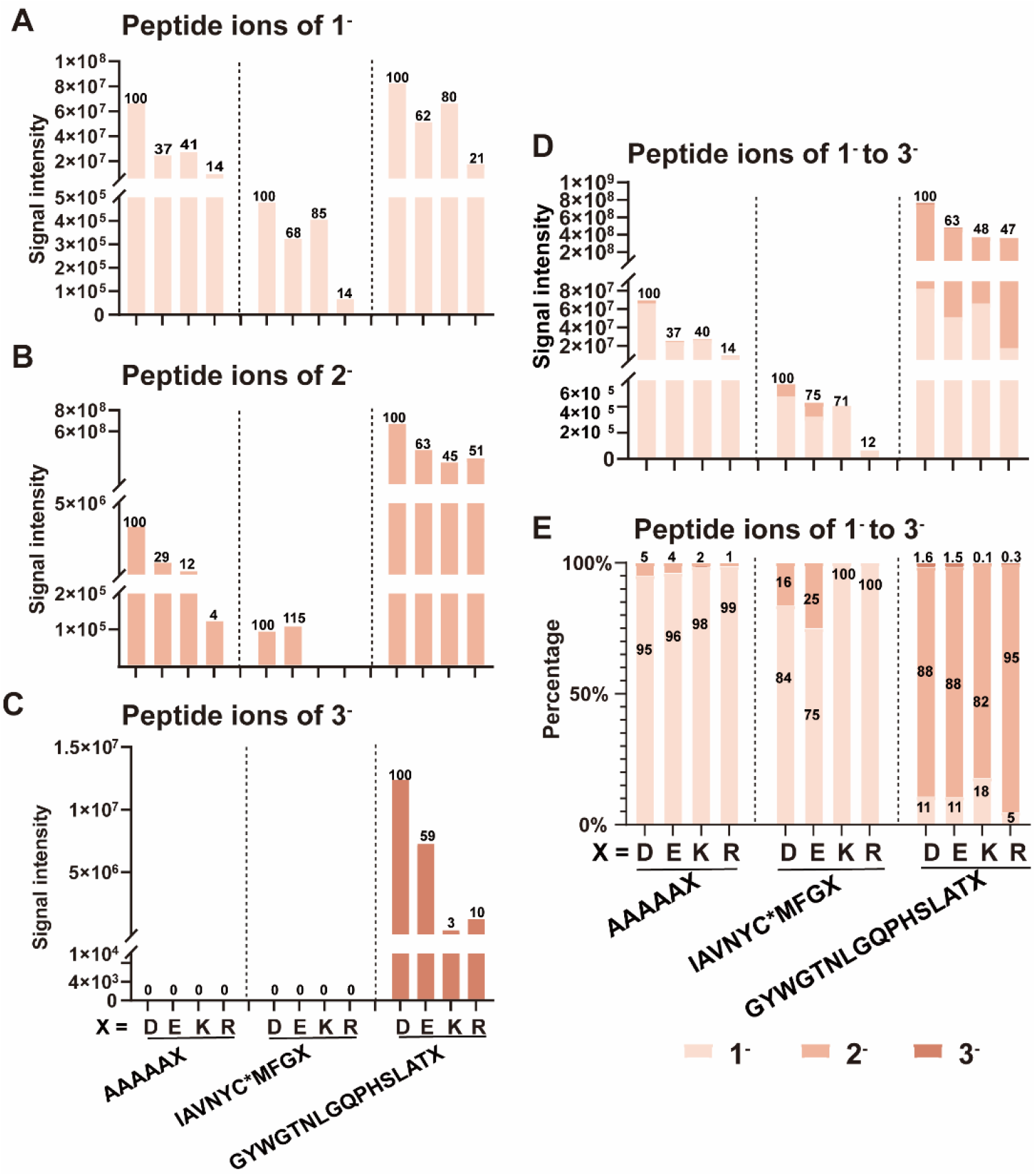
MS1 intensity of 12 synthetic peptides normalized against the relative concentration shown in Table 1. MS1 intensity of 1^-^ (A), 2^-^ (B), 3^-^ (C), and 1^-^ to 3^-^ (D) precursors. (E) Relative MS1 intensities of 1^-^, 2^-^, and 3^-^ precursors. The numbers indicate the relative intensity in percentage. *Carbamidomethylated cysteine.

**Figure 2.**
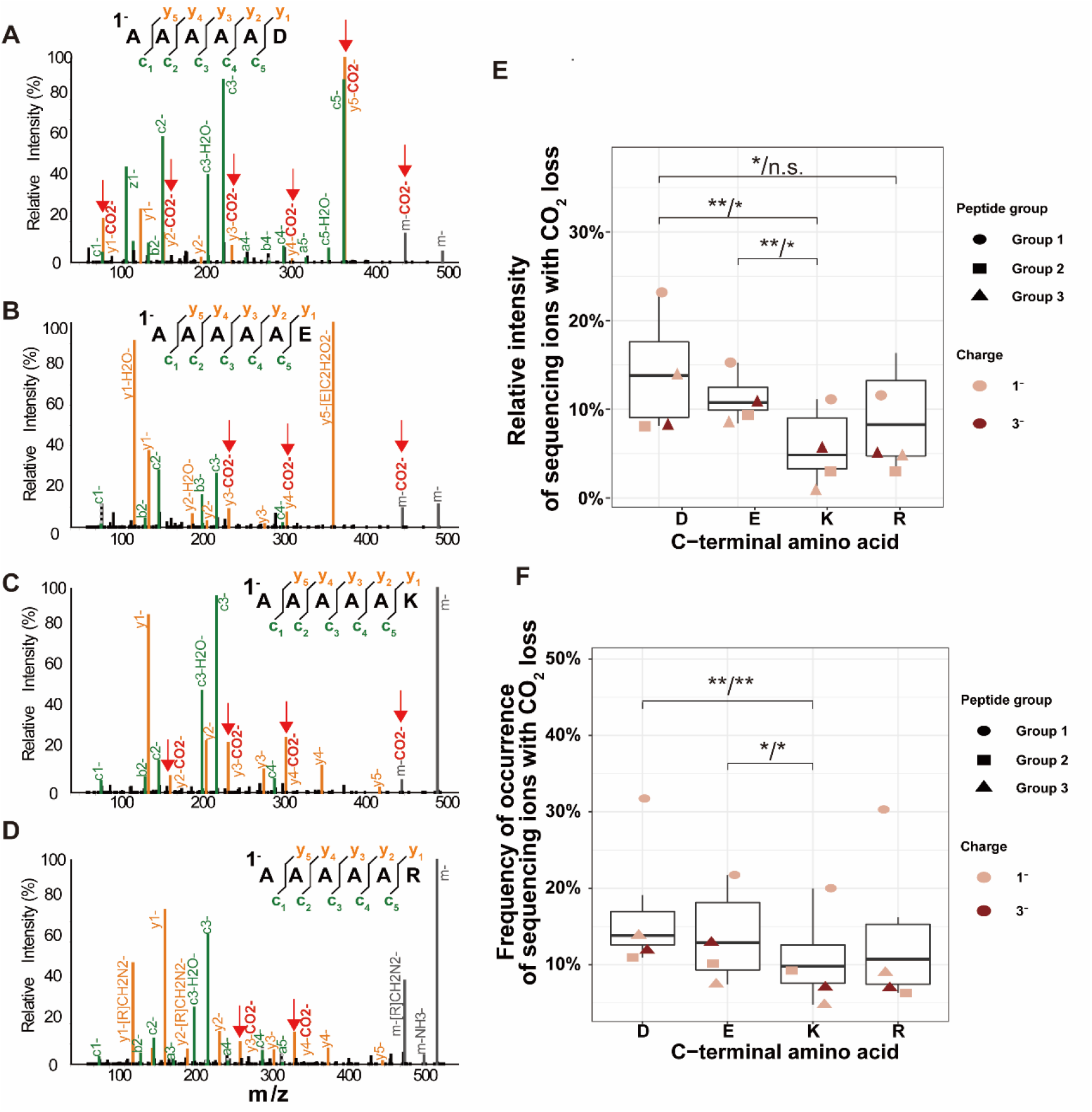
Frequent CO_2_ loss in HCD spectra of deprotonated peptides. HCD spectra of AAAAAD (A), AAAAAE (B), AAAAAK (C), and AAAAAR (D) at NCE 35. The peaks matched to theoretical peptide fragments within mass tolerance of 10 ppm were labeled, and matched ions with CO_2_ loss are highlighted and indicated by red arrows. The original high resolution spectra are showed in Figure S1. The relative intensity (E) and frequency of occurrence (F) of sequencing ions with CO_2_ loss for peptides ending in D, E, K, or R. The relative intensity was calculated by dividing the summed intentisities of sequencing ions with CO_2_ loss with the summed intentisities of all ions found in a MS2 spectrum, and expressed in percentage. The frequency of occurrence was calculated by dividing the number of sequencing ions with CO_2_ loss with the total number of ions in a MS2 spectrum, also expressed in percentage. Paired t-test and nonpaired Wilcoxson rank-sum test were used to compare between groups. Both *p* values are labeled on the plot (t-test/Wilcoxson test) if either is less than 0.05; n.s. *p* ≥ 0.05; * *p* < 0.05; ** *p* < 0.01. One representative HCD spectrum was used for each charge state of each peptide. Among the group 2 peptides, only the ones with C-terminal D or E had 2^-^ precursors, so the two corresponding HCD spectra were not used in the paired t-test when comparing either D or E to K or R.

**Figure 3.**
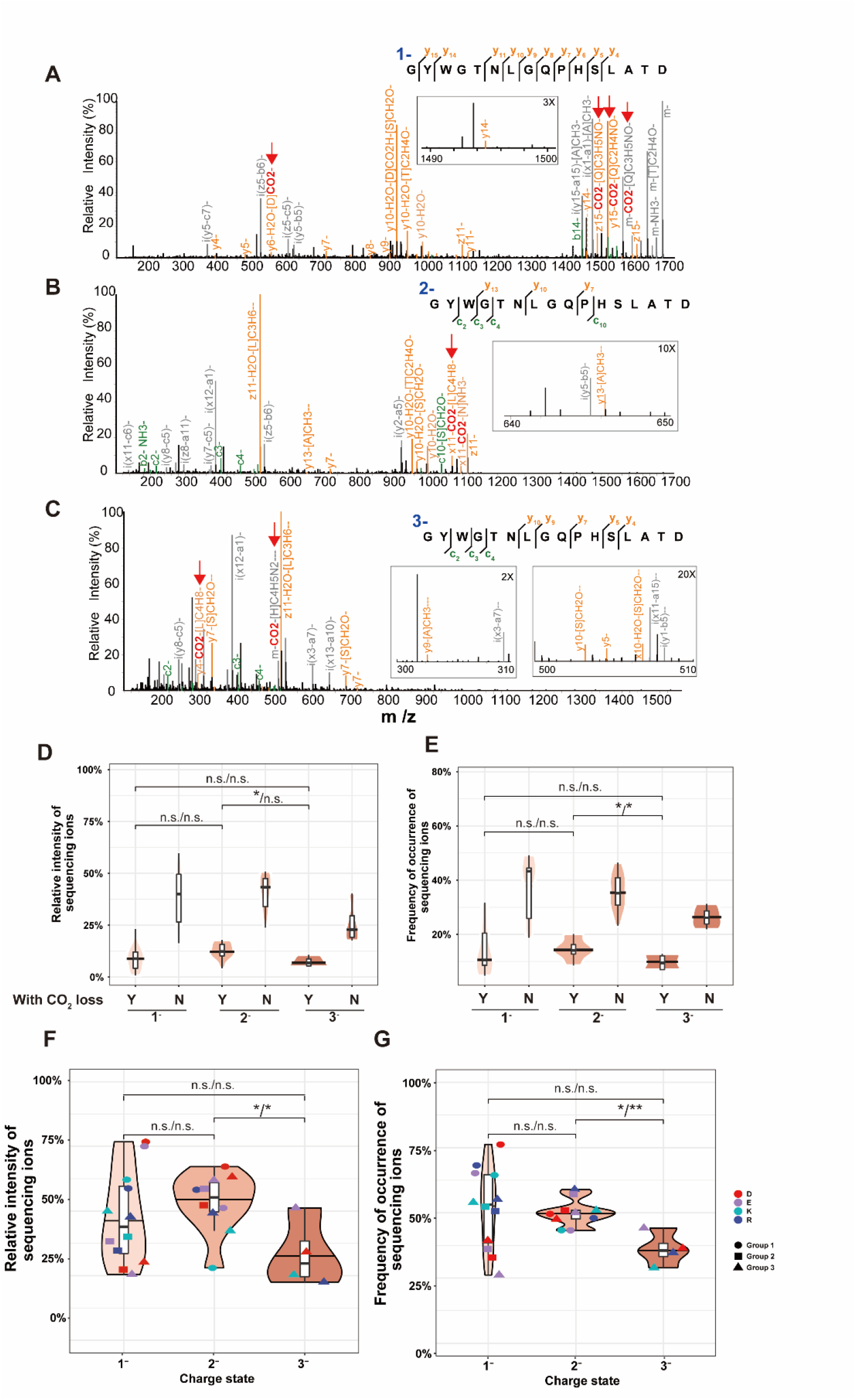
Neutral loss of CO_2_ and the amount of sequencing ions in the HCD spectra of 1^-^, 2^-^, and 3^-^ peptide precursors. HCD spectra of GYWGTNLGQPHSLATD from a 1^-^ (A), 2^-^ (B), or 3^-^ (C) precursor. The peaks matched to theoretical peptide fragments within mass tolerance of 10 ppm were labeled, and matched ions with CO_2_ loss are highlighted and indicated by red arrows. The original high resolution spectra are showed in Figure S2. Relative intensity (D) and frequency of occurrence (E) of sequencing ions that had a neutral loss of CO_2_ loss were compared across precursor charge states using nonpaired t-test/nonpaired Wilcoxon rank sum test. Relative intensities (F) and frequency of occurrence (G) of all sequencing ions were compared across precursor charge states using paired t-test/nonpaired Wilcoxon rank sum test. ** *p* < 0.01;* *p* < 0.05; n.s. *p* ≥ 0.05. The relative intensity of sequencing ions was calculated by dividing the summed intentisities of sequencing ions with the summed intentisities of all ions found in a MS2 spectrum. The frequency of occurrence was calculated by dividing the number of sequencing ions with the total number of ions in a MS2 spectrum.

An additional collection of 115 synthetic peptides as listed in Supplemental Table S1 were used in this study. Among them, 1-37 were used earlier for a different purpose (*25*), 38-55 were provided by Dr. Ping Xu from National Center for Protein Sciences (Beijing, China), and 56-115 were synthesized by Scilight Biotechnology (Beijing, China). Each peptide was brought to 10 μM in 7.5 mM hexylamine, 50% acetonitrile before being sprayed into mass spectrometer as previously reported (*23*).

### 2.2 MS parameters

All experiments were performed on Orbitrap Fusion™ Lumos™ Tribrid™Mass Spectrometer (Thermo Fisher Scientific). For the synthetic peptides, the sample solution was introduced into the electrospray source at a flow rate of 2 μL/min with a syringe pump. The ion spray voltage was 2.5 kV in the negative ion mode. The temperature of ion transfer tube was 320 °C. The MS1 mass ranges were 150 to 2000 m/z and adjusted according to the mass of each peptide, while the MS2 mass ranges were adjusted automatically. Each MS1 or MS2 spectrum was an averaged spectrum produced using 3 microscans. The AGC (Automatic Gain Control) target was 1E6 for MS1 and 1E5 for MS2, maximum injection time was adjusted automatically both for MS1 and MS2. The resolution was 120K for both MS1 and MS2. The inclusion list contained the theoretical m/z values of each peptide in the sample—from [M-H]^-^ to [M-9H]^9-^. After a precursor was detected with a mass tolerance of 20 ppm, they were selected for fragmentation using an isolated window of 0.4 m/z for the 4 peptides of group 2 and 2.0 m/z for all other peptides. For each precursor, HCD spectra were acquired with varied normalized collision energy (NCE) from 10 to 50, with a step length of 5. MS1 and MS2 spectra were recorded, respectively, in the profile mode and the centroid mode.

### 2.3 Data analysis

The HCD spectra acquired at the optimal NCE of 35 were analyzed using in-house Python scripts. Theoretical *m/z* values of the precursors, the a-, b-, c-, x-, y-, z-fragment ions and their neutral loss products, and the internal ions caused by double cleavage of the precursors were calculated and compared with the observed peaks on MS2 spectra. Abundant neutral losses were taken into consideration, including loss of H_2_O, NH_3_, or CO_2_ loss, which are not specific for a particular amino acid, and loss of characteristic amino acid side chains. In total, 66 amino acid side chain neutral losses (NL) were taken into consideration, 65 were reported by Bowie et al. (*11*), Liang et al. (*18*), Rumachik et al. (*19*), and Sun et al.(*26*), and the other one was found in this study (58.005479, C_2_H_2_O_2_ from E) (Supplemental Table S2-3). The reported NLs from oxidized methionine (*18*) were not examined. For each MS2 spectrum of a given peptide, the experimental peaks were matched to the theoretical ions with a mass tolerance of 10 ppm. Of the experimental peaks, only those with a relative intensity (intensity of peak/base peak intensity) above 0.01 were included for statistical analysis unless indicated otherwise. Protein isoelectric points were calculated using http://isoelectric.org/ from input sequence files.

## 3. Results

We started out by investigating how an acidic C-terminus (D/E) generated by Glu-C digestion affects MS analysis of deprotonated peptides compared to a basic C-terminus (lysine/arginine, or K/R) generated by trypsin digestion. It has been reported that fragmentation of Glu-C peptides seems to be more extensive than tryptic peptides in NETD analysis (*6*). We thus wondered whether Glu-C digestion might be advantageous over trypsin digestion for negative HCD analysis of acidic proteins. A total of 12 synthetic peptides in 3 groups (Table 1) were analyzed. Within each group, the four peptides differ only in the C-terminal residue, which is either D, E, K, or R. The length of these peptides varied from 6 to 16 residues, and the 20 common amino acids found in proteins were all included in these peptides.

An equimolar mixture of the four peptides of the same group was sprayed directly into an Orbitrap Fusion Lumos mass spectrometer operated under the negative ion mode. MS1 and HCD MS2 spectra were acquired. For HCD, the optimal normalized collision energy (NCE) of 35 was used (*23*). Figure 1 shows a summary of the data. Group 1 (AAAAAX) and group 2 (IAVNYC*MFGX) peptides, in which X is D, E, K, or R, generated predominantly 1^-^ ([M-H]^-^) precursor ions and no 3^-^ ([M-3H]^3-^) precursors. The group 3 peptides (GYWGTNLGQPHSLATX) generated predominantly 2^-^ ([M-2H]^2-^) precursors and small amounts of 1^-^ and 3^-^ precursors (Figure 1A-C). Within each group, the peptide ending in D had the highest total intensity in MS1 whereas the peptide ending in R had the lowest (Figure 1D). The sequence-matched E/K peptides, i.e., two peptides differed only in the C-terminal residue—either E or K, displayed no obvious difference in total MS1 intensity (Figure 1D). As expected, for each of the three groups of peptides, the presence of a single D or E consistently shifted the charge state distribution of the [M-nH]^n-^ ions upwards (Figure 1E).

The negative-ion HCD spectra of the 12 peptides displayed widespread NL of CO_2_, regardless of the identity of the C-terminal aa (Figure 2). We observed loss of CO_2_ off precursor ions or fragment ions, which include those that inform the sequence of the peptide (*a-, b-, c-, x-, y-*, or *z-*, produced after the peptide backbone is cleaved once at any position) and thus referred to as sequencing ions (Figure 2). With respect to the relative intensity of sequencing ions that have lost one or more CO_2_ molecules, the peptides with a C-terminal D/E tend to be higher than their K/R counterparts (Figure 2A-E). Those with a C-terminal K produced the least amount of CO_2_ loss (Figure 2E-F). Pairwise comparison finds that the difference is statistically significant (*p* < 0.05, paired *t*-test) between D and K, E and K, and D and R (Figure 2E). Regarding the frequency of occurrence of sequencing ions with CO_2_ loss, the peptides with a C-terminal D or E are significantly higher than those with a C-terminal K (Figure 2F, *p* < 0.05, paired *t*-test).

Peptide anions of different charge states all produce CO_2_-loss peaks in HCD spectra. Shown as an example in Figure 3 are the HCD spectra of 1^-^, 2^-^, and 3^-^ precursors of the same peptide (Figure 3A-C). Across the three charge states, the sequencing ions that had a neutral loss of CO_2_ accounted for, in most cases, less than 20% of all sequencing ions in terms of peak intensity (Figure 3D, expressed as relative intensity) or peak number (Figure 3E, expressed as frequency of occurrence).

In an earlier study of 36 peptides, we observed that negative-ion HCD at NCE of 35, which was the global optimum, produced higher-intensity sequencing ions from 1^-^ and 2^-^ precursors than from 3^-^ precursors (*23*). Analysis of the negative-ion HCD spectra of the sequence-matched peptides (Table 1), also acquired at NCE 35, confirmed that 2^-^ precursors had higher-intensity and a greater number of sequencing ions than 3^-^ precursors (Figure 3F-G, *p* < 0.05, paired *t*-test). FigureThe difference between 1^-^ and 3^-^ precursors was not significant in this 12-peptide dataset.

The pervasiveness of CO_2_ loss was to some extent beyond our expectation. Loss of CO_2_ from deprotonated peptides upon CID had been reported (*5, 7, 16, 18, 27*), but it was ranked at the 10^th^ (from glutamic acid) or 14^th^ (from aspartic acid) position by relative abundance (*18*). In our previous analysis (*23*), only the reported eight most abundant NLs from aa side chains (*18*) were taken into consideration. Prompted by the unexpectedly extensive loss of CO_2_ in the HCD spectra of deprotonated peptides (Figure 1-3), we repeated the statistical analysis of NL peaks by considering all of the reported NL species (*11, 18, 19, 26*) on an extended dataset of 115 synthetic peptides of 5-28 aa (Supplemental Table S1), which included 21 previously analyzed peptides (*23*) but none of the peptides in Table 1. Table 2 shows a summary of the analysis result. The most abundant NL species include a loss of NH_3_ from asparagine (N), a loss of C_2_H_4_O from threonine (T), and a loss of H_2_O or CO_2_ from D and E. We also found a previously unnoticed NL of 58.005479 (C_2_H_2_O_2_) from E (Table 2 and Figure 2B).

**Table 2.**
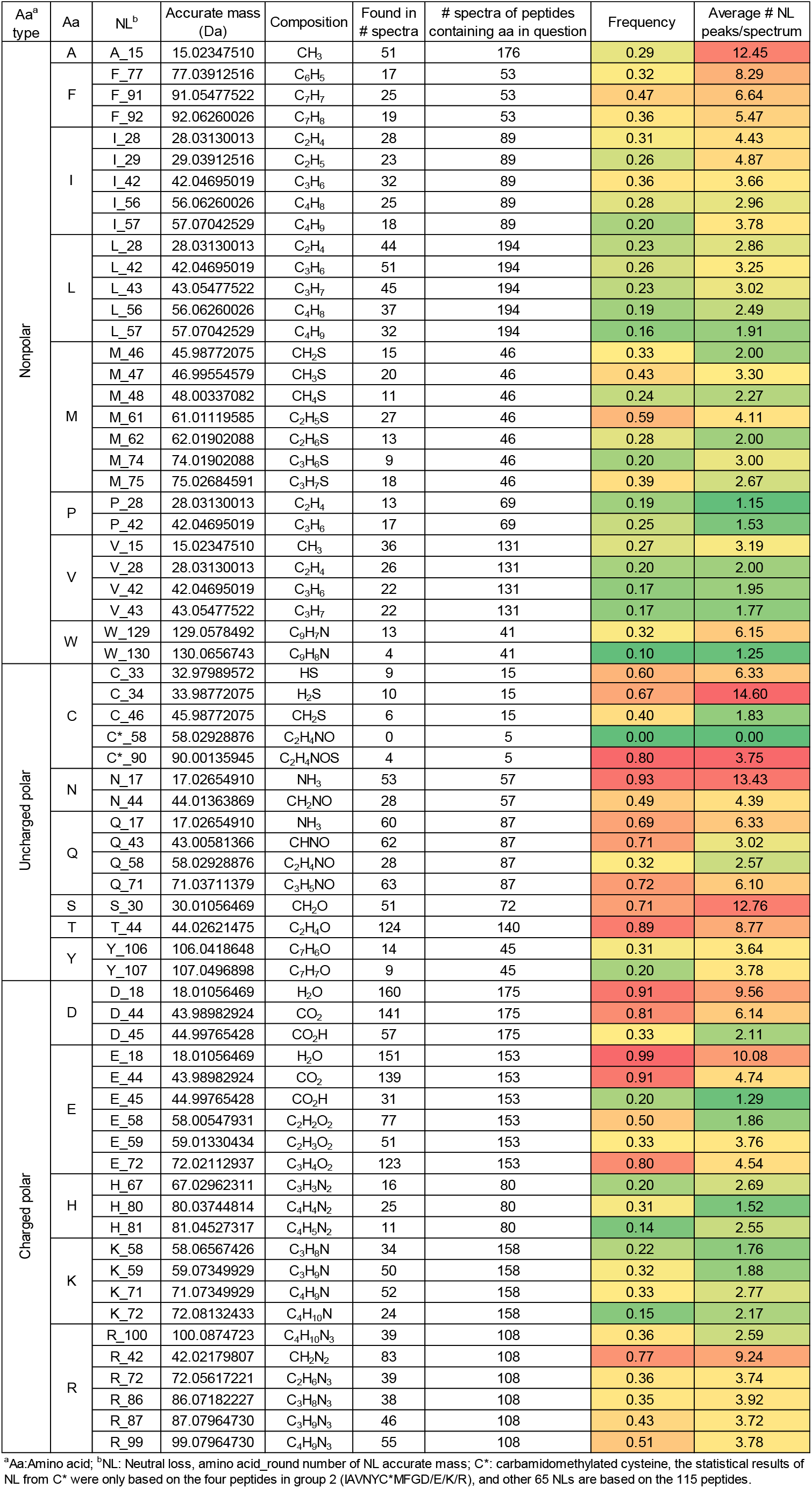
Side-chain neutral losses observed from 115 peptides

The relative abundance of the *a-, b-, c-, x-, y-*, and *z-* ions with or without various types of NLs is updated (Figure 4). As loss of CO_2_ does not require the presence of D or E in a peptide (Figure 2C-F), it is regarded as a common NL, although it is most strongly associated with D and E. The updated result shows again that the majority of the fragment ions have NLs of one form or another, including any of the common NLs (H_2_O, NH_3_, CO_2_) and aa-specific side-chain losses. The four most abundant fragment ion types are *y-* > *x-, z-* > *c-* if the NL ions are included, and *c-* > *y-* > *z-* > *b-* if not(Figure 4).

**Figure 4.**
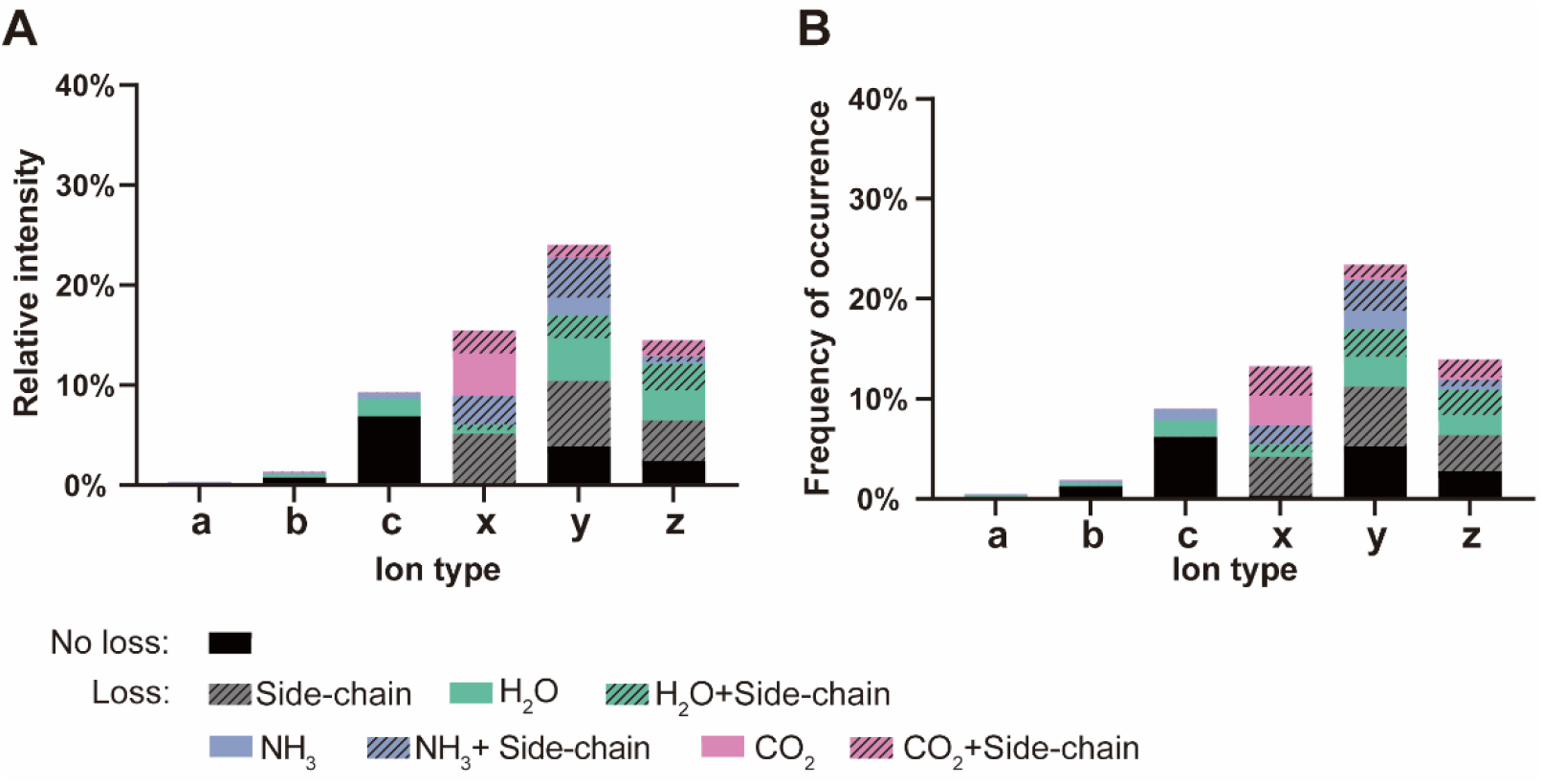
Relative intensity (A) and frequency of occurrence (B) based on 254 HCD spectra of 115 synthetic peptides. Note: y-H_2_O and x-CO_2_ have the same mass. As a result, the peaks matched to the two ion types were counted twice.

The dataset of 115 synthetic peptides allowed a charge-state distribution analysis. As shownFigure, longer peptides and D/E-rich peptides are more likely to carry multiple negative charges (Figure 5). The peptides with <15 aa and no more than 3 D/E residues produced predominantly [M-H]^-^ and [M-2H]^2-^ ions, whereas the peptides containing 4-7 D/E residues carried up to five negative charges.

**Figure 5.**
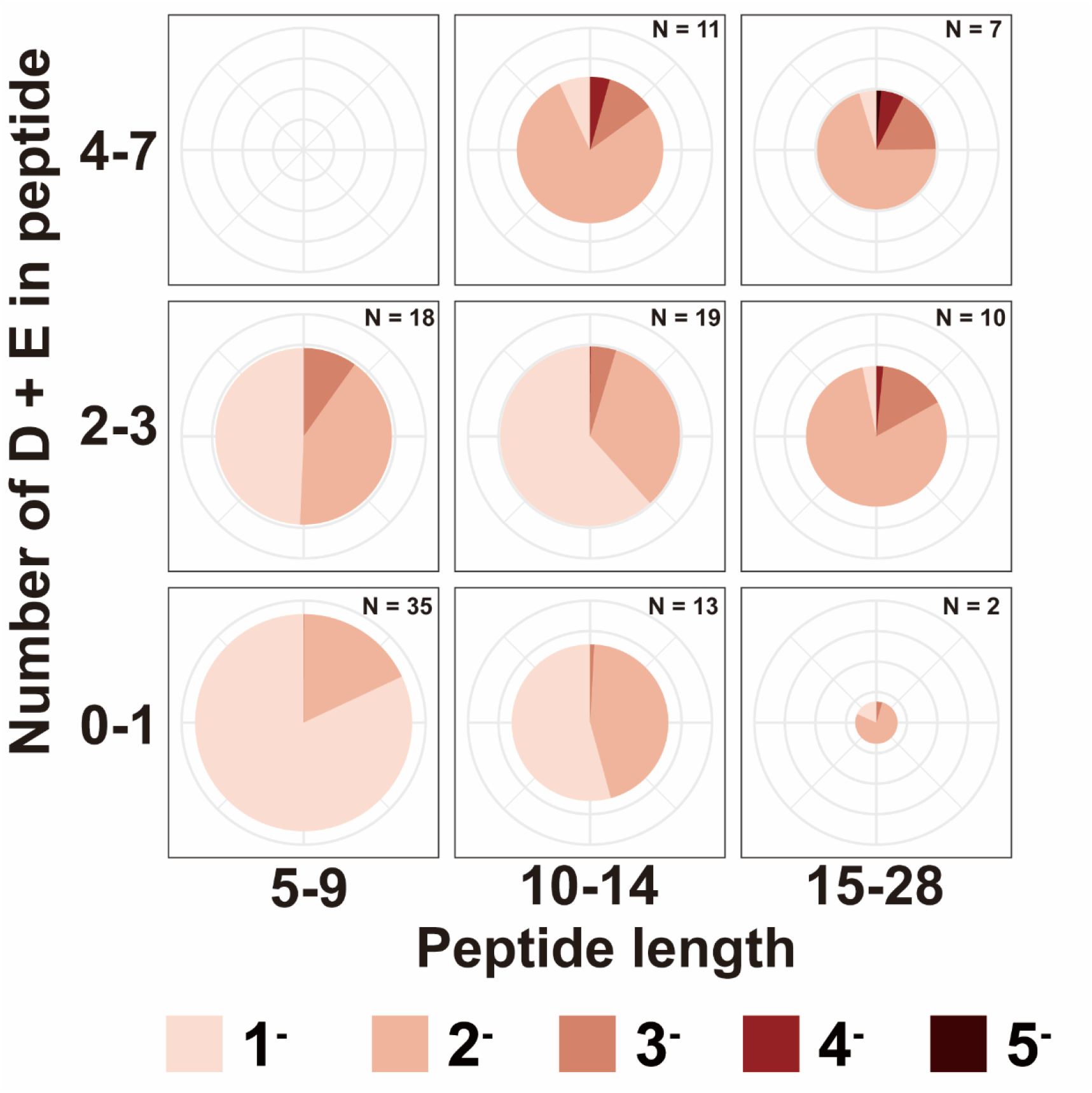
The length and the number of acidic residues of a peptide influence the charge state of the peptide anion. Each circle represents a group of peptides of the indicated length and the indicated number of D/E residues. The diameter of a circle reflects the number of peptides in a group. The sectors within a circle each represent a charge state observed in MS1, and the area of a sector indicates the average occupancy of an of ions of the indicated charge state. This statistics is based on the MS1 spectra of 115 synthetic peptides. The precursor ions were identified from MS1 using a mass tolerance window of 20 ppm centered around the calculated m/z values.

Using this expanded dataset, we analyzed the effect of D/E residues on the HCD spectra of deprotonated peptides. As shown, D/E-containing peptides produced more and higher-intensity CO_2_-loss peaks than D/E-free peptides (Figure 6A-B). Further, the subgroup of peptides containing only C-terminal D/E may tend to lose CO_2_ more readily than the subgroup of peptides containing only non-C-terminal D/E (Figure 6C-D). The difference is significant in terms of frequency of occurrence (Figure 6D), not in relative intensity (Figure 6C).

**Figure 6.**
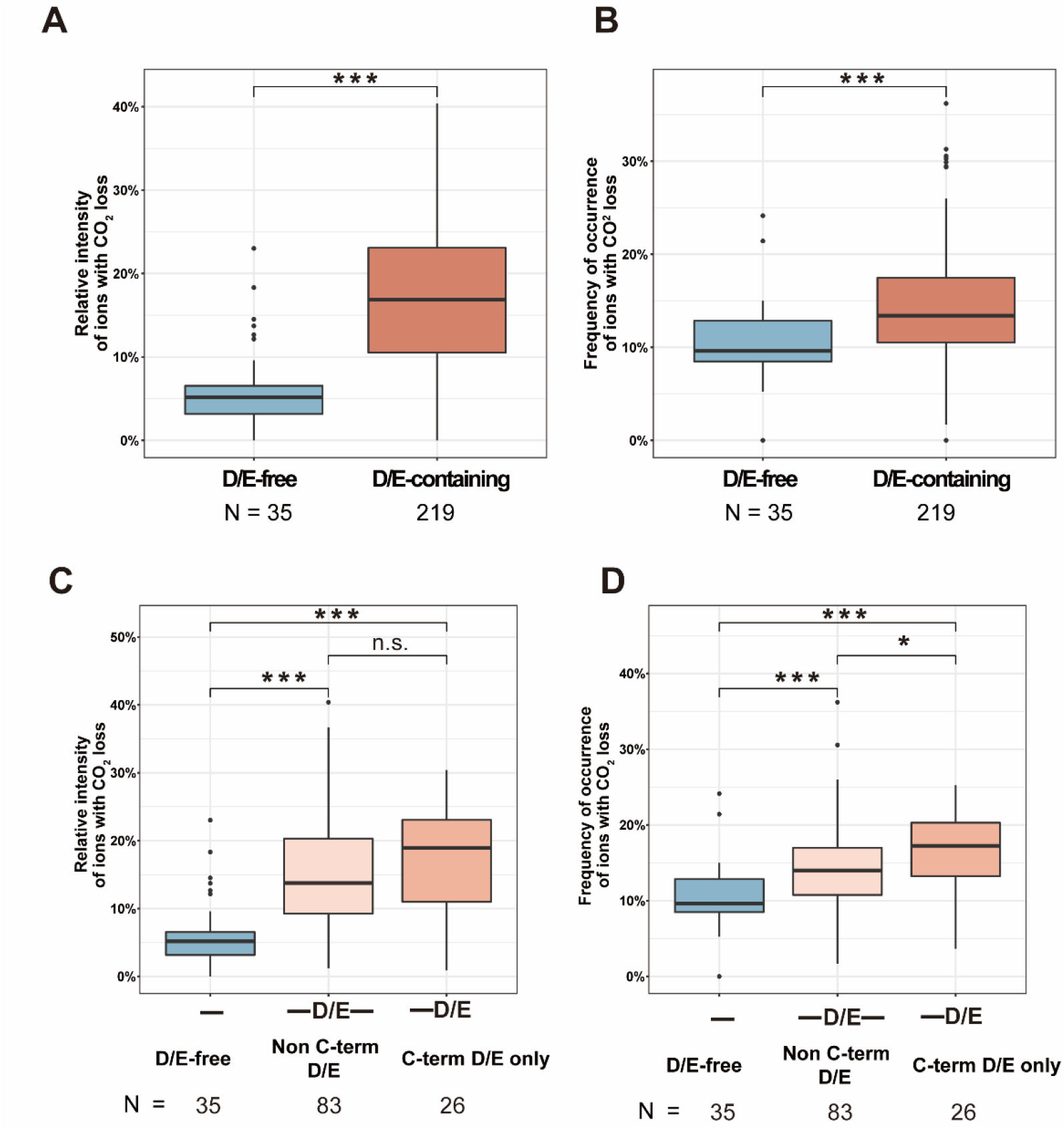
Peptides containing D/E residues have more neutral loss of CO_2_ in negative-ion HCD. Relative intensities (A, C) and frequency of occurrence (B, D) of any ions with CO_2_ loss, based on 254 HCD spectra of 115 peptides. The relative intensity here was calculated by dividing the summed intentisities of any ions with CO_2_ loss with the summed intentisities of all ions found in a MS2 spectrum. The frequency of occurrence was calculated by dividing the number of any CO_2_-loss ions with the total number of ions in a MS2 spectrum. (A, B) D/E-free peptides versus D/E-containing peptides. (C, D) Comparion between D/E-free peptides, peptides containing non C-terminal D/E residues, and peptides containing only C-termimal D/E residues. *** *p* < 0.001;** *p* < 0.01; * *p* < 0.05; n.s. *p* ≥ 0.05; nonpaired Wilcoxson rank-sum test.

Similar to the result obtained from sequence-matched peptides (Figure 3D-E), the negative-ion HCD spectra of 115 peptides showed that sequencing ions had more or less similar levels of CO_2_ loss across precursor charge states from 1^-^ to 4^-^ (Figure 7A-B). Examining the sequencing ions as a whole, we found that the percentage of sequencing ions in the HCD spectra of the [M-H]^-^ and [M-2H]^2-^ precursors tend to be higher than that of the [M-3H]^3-^ and [M-4H]^4-^ precursors (Figure 7C-D). This is in agreement with the finding of an earlyier study (*23*) and that from sequence-matched peptides (Figure 3F-G). This suggests that peptide sequencing by negative-ion-mode MS should focus mainly on lower-charge state peptide ions. Long peptides and D/E-rich peptides tend to produce higher charge state anions (Figure 5), we thus propose that relatively short peptides containing no more than a couple of D/E residues are potentially suitable for MS sequencing in the negative ion mode.

**Figure 7.**
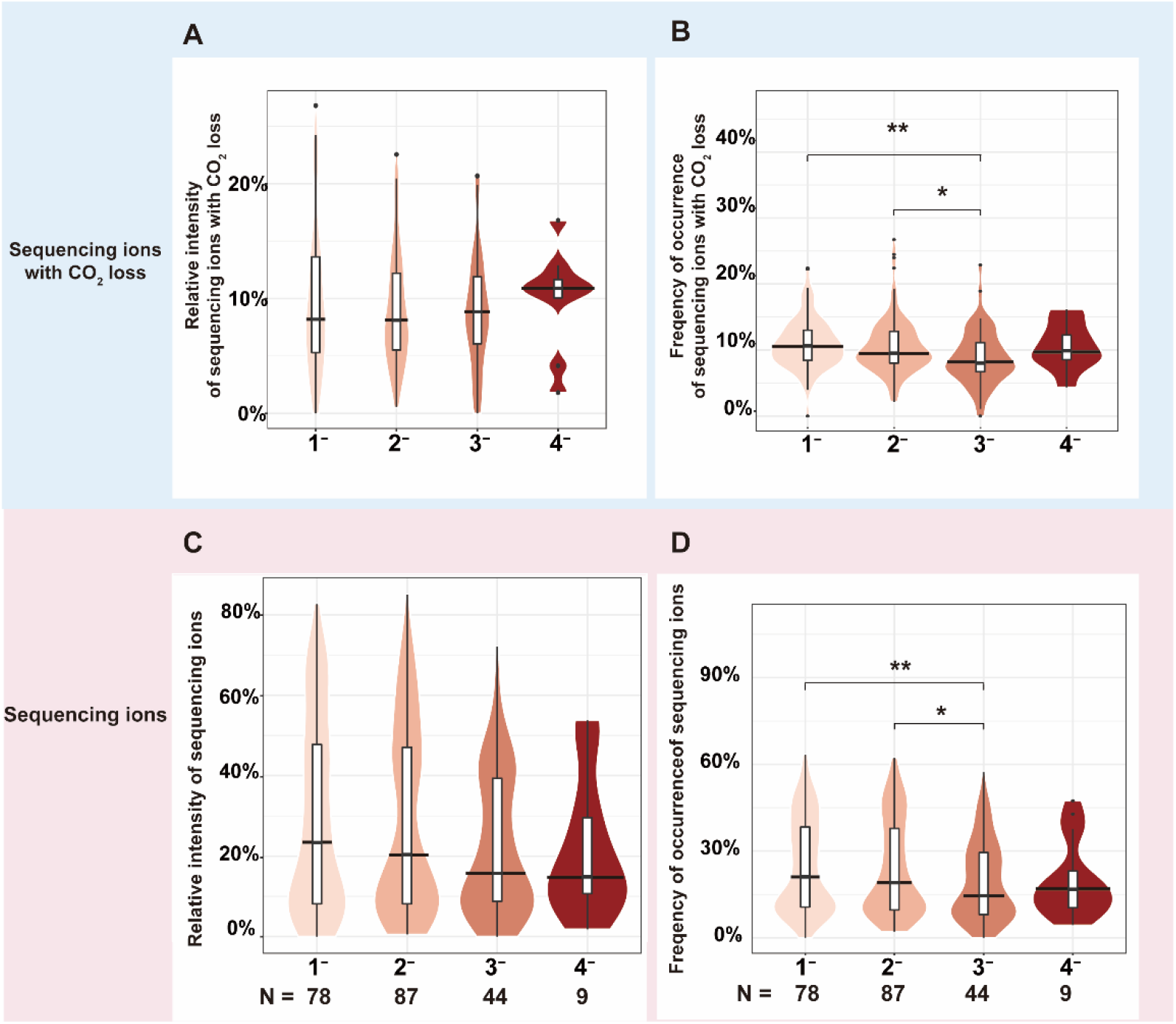
Influence of precursor charge state on sequencing ions. Relative intensities (A, C) and frequency of occurrence (B, D) of sequencing ions with CO2 loss (A, B) or all sequencing ions as a whole (C, D), based on 254 negative-ion HCD spectra of 115 peptides. The relative intensities and the frequency of occurrence were calculated as described in Figure 1 and Figure 2. Nonpaired Wilcoxson rank-sum test, *p* values less than 0.1 are labeled, ** *p* < 0.01; * *p* < 0.05.

With this knowledge, we performed in silico Glu-C or trypsin digestion of a standard protein bovine serum albumin (BSA, calculated isoelectric point 5.59) and a highly acidic protein pepsin (calculated isoelectric point 3.86). Pepsin, itself a protease active in acidic environments, is a 385-aa protein with 45 D/E (11.7%) and 15 K/R (3.9%) residues (Supplemental Figure 3A-B). BSA is a 607-aa protein with 99 D/E (16.3%) and 86 K/R (14.2%) residues (Supplemental Figure 3C-D). Allowing no more than two missed cleavage sites, in silico digestion of BSA by trypsin or Glu-C generated 137 or 196 peptides, respectively, of less than 20 aa and no more than 3 D/E residues, which are potentially suitably for sequence identification by negative-ion HCD (Figure 8A-B). This suggests that either protease can be used to digest BSA, albeit trypsin might be a better one. In contrast, Glu-C is clearly a better choice for pepsin since in silico digestion of pepsin by trypsin produced only 25 peptides potentially suitable for negative-ion HCD, that by Glu-C produced 69 (Figure 8C-D). This lends support to the idea that Glu-C digestion is worth trying for negative-ion mode MS analysis of highly acidic proteins.

**Figure 8.**
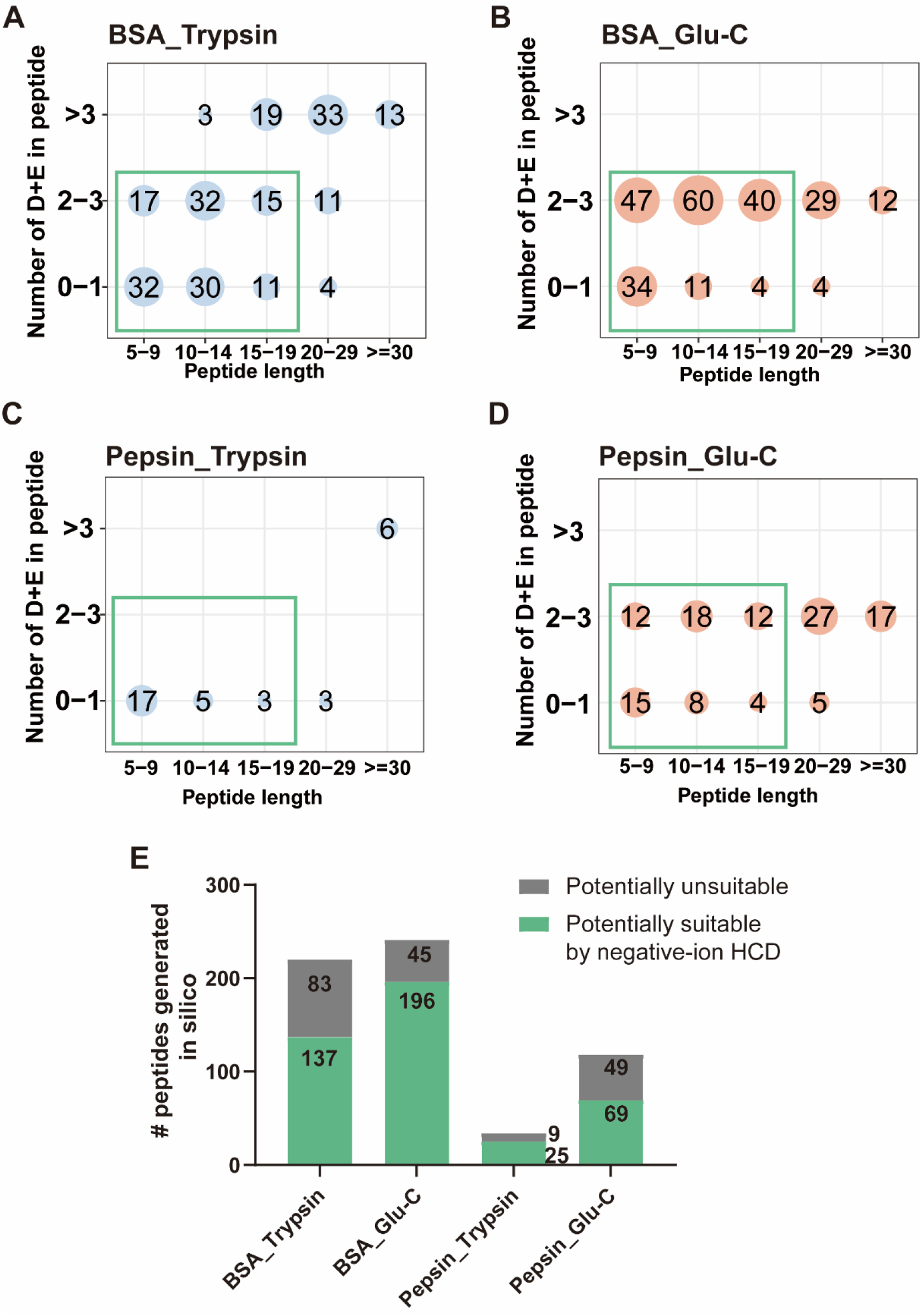
In silico Glu-C digestion of highly acidic proteins generate more peptides that are potentially suitable for negative-ion HCD. Peptides from in silico digestion of BSA (A, B) and pepsin (C, D) with trypsin (A, C) or Glu-C (B, D). The number in each circle indicates the number of peptides of the indicated length and containing the indicated number of acidic residues. Peptides potentially suitable for sequencing by negative-ion HCD are framed by a green rectangle.

## 4. Discussion

Many types of NLs disperse the signal of a fragment ion into different channels and thus decrease the intensity of fragment ions. This reduces the sensitivity of peptide identification in negative-ion-mode MS. How to suppress NL is worth investigating in the future. Methylation of side-chain (D/E) and the C-terminal -COOH is one possibility to reduce neutral loss of CO_2_.

Although Glu-C can cleave the peptide bond C-terminal to Asp (*6, 28, 29*), the efficiency of this cleavage is lower than its namesake cleavage—cutting after a Glu residue. Another protease Asp-N cleaves the peptide bond N-terminal to D. The frequency of D, E, K, and R in the 567483 protein sequences of the UniProtKB database is 5.46%, 6.72%, 5.80%, and 5.53%, respectively (UniProtKB/Swiss-Prot protein knowledgebase release 2022_02 statistics, https://web.expasy.org/docs/relnotes/relstat.html). So, the frequency of cleavage by Asp-N and Glu-C combined (12.18%) and that by trypsin (11.33%) are similar, which means that they will cleave proteins into peptides of similar length distributions. The difference lies in the number of D/E residues, which will vary greatly in tryptic peptides but remain low in peptides generated by Asp-N and Glu-C double digestion. Peptide anions containing multiple D/E residues tend to adopt higher charge states, and higher charge state precursors (e.g. 3^-^ or above) produce poorer MS2 spectra with lower-intensity sequencing ions (Figure 7). Therefore, protein digestion with Asp-N and Glu-C together may be a worthy alternative for negative-ion-mode MS analysis.

The statistical analysis in this study did not look into locally enhanced backbone cleavage by specific aa side chains, such as the cleavage at the N-Cα bond of X-D/N or X-S/T, which has been reported (*7, 11, 22, 25*). Certain amino acid residues including aspartic acid, glutamic acid, and cysteine have been shown to influence locally the backbone cleavage of deprotonated peptides (*8, 11, 12, 17, 22, 30*). This waits further analysis.

Previously, we marveled at the observation that the highly basic peptides containing multiple lysine residues, zero acidic residues, and amidated C-termini ionized well in the negative ion mode and carried up to three negative charges (23). One example is a 16-aa peptide GLFAVIKKVAKVIKKL-NH2, which was first studied by Dr. Bowie (11-13). In this study, our side-by-side comparison of the peptides differing only in the C-terminal residue (Table 1) showed that replacing the C-terminal E with K did not decrease the total intensity of the deprotonated peptides (Figure 1D), suggesting that lysine has little negative effect on peptide ionization in the negative ion mode. A K-rich and D/E-free peptide, if sufficiently long, is able to carry multiple negative charges (Figure 5). Further, their 1^-^ and 2^-^ precursors can generate beautiful MS2 spectra with abundant sequencing ions (Figure 7). This is most noticeable if they are devoid of other NL-prone aa residue besides D and E, for example, C, N, Q, S, T, and R (Table 2). The peptide GLFAVIKKVAKVIKKL-NH2 happens to have all the desirable attributes.

In summary, we have found evidence to suggest that contrary to what might be expected, high acidic peptides with multiple D/E residues should be difficult to identify by negative-ion HCD. This is because D/E-rich peptides tend to form multiply charged anions, whose HCD spectra generally have a scarcity of sequencing ions. Peptide anions of 1^-^ or 2^-^ charge state may be more suitable for sequence identification by HCD because they produce a higher fraction of sequencing ions than multiply deprotonated peptides. Loss of CO_2_, although common to all deprotonated peptides regardless of their charge state, is enhanced in the HCD spectra of D/E-containing peptides. Additionally, we found that in the HCD spectra of peptide anions, the most abundant fragment ion types are y-> x-, z- > c- if the NL ions are included, or c- > y- > z- > b- if not. These results will inform future studies aiming to uncover the fragmentation mechanisms of deprotonated peptides, as well as method development for peptide sequencing by negative-ion HCD.

## Acknowledgments

The authors would like to thank Dr. Ping Xu of the National Center for Protein Sciences (Beijing, China) for providing 18 synthetic peptides, and Peng-Zhi Mao of the Institute of Computing Technology, Chinese Academy of Sciences for modifying pLable software to display peptide anion spectra, and Jishuai Zhang of National Institute of Biological Sciences, Beijing, for preparation of the graphic abstract. This study is supported by National Key Research and Development Program of China (Grant 2020YFF01014505), the Ministry of Science and Technology of China, and Beijing Municipal Science and Technology Commission.

---

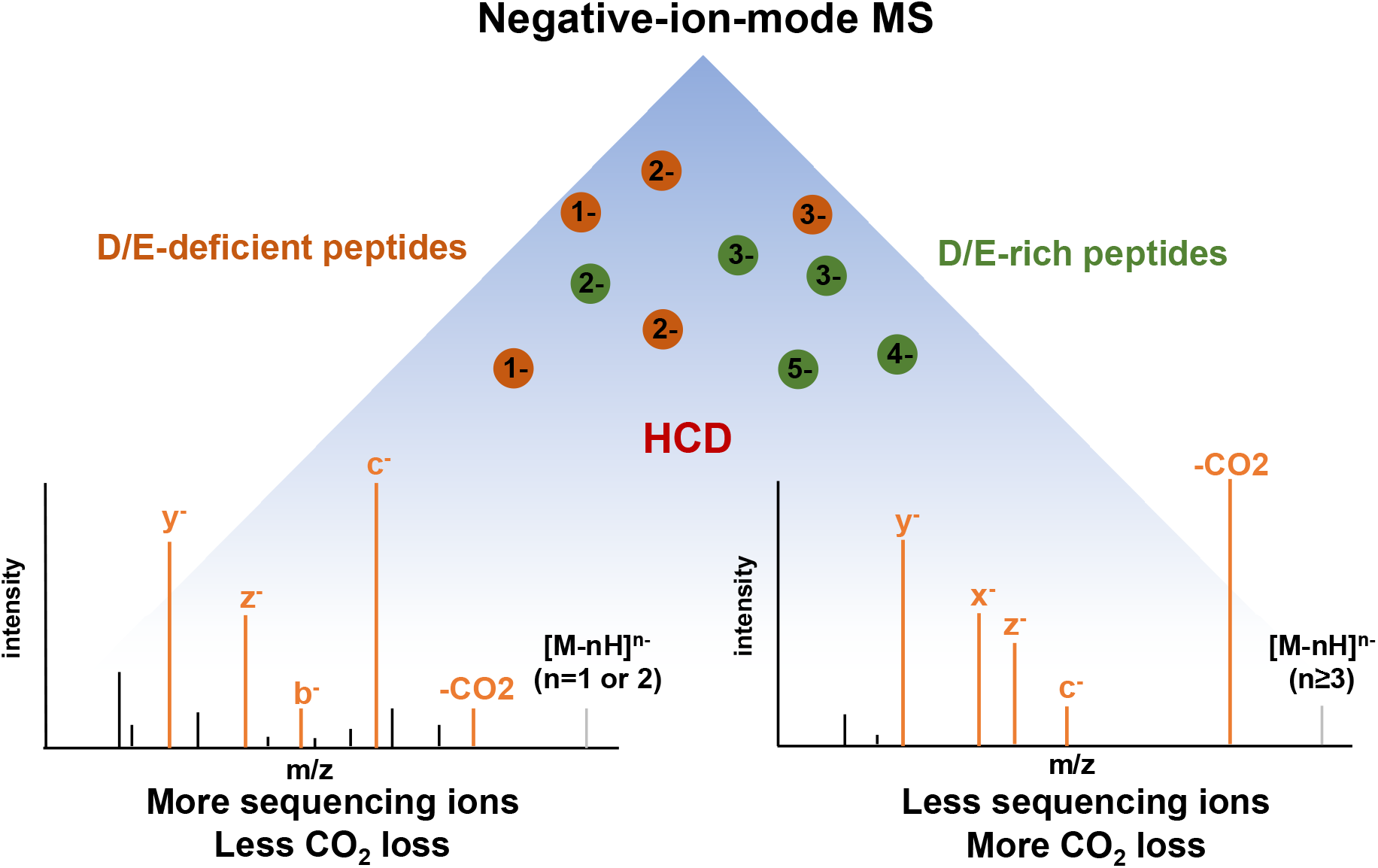

