## Supplement Fig 1-3 for "D/E-rich peptides are less suitable than D/E-deficient peptides for identification by negative-ion HCD due to scarce production of sequencing ions from multiply charged precursors"

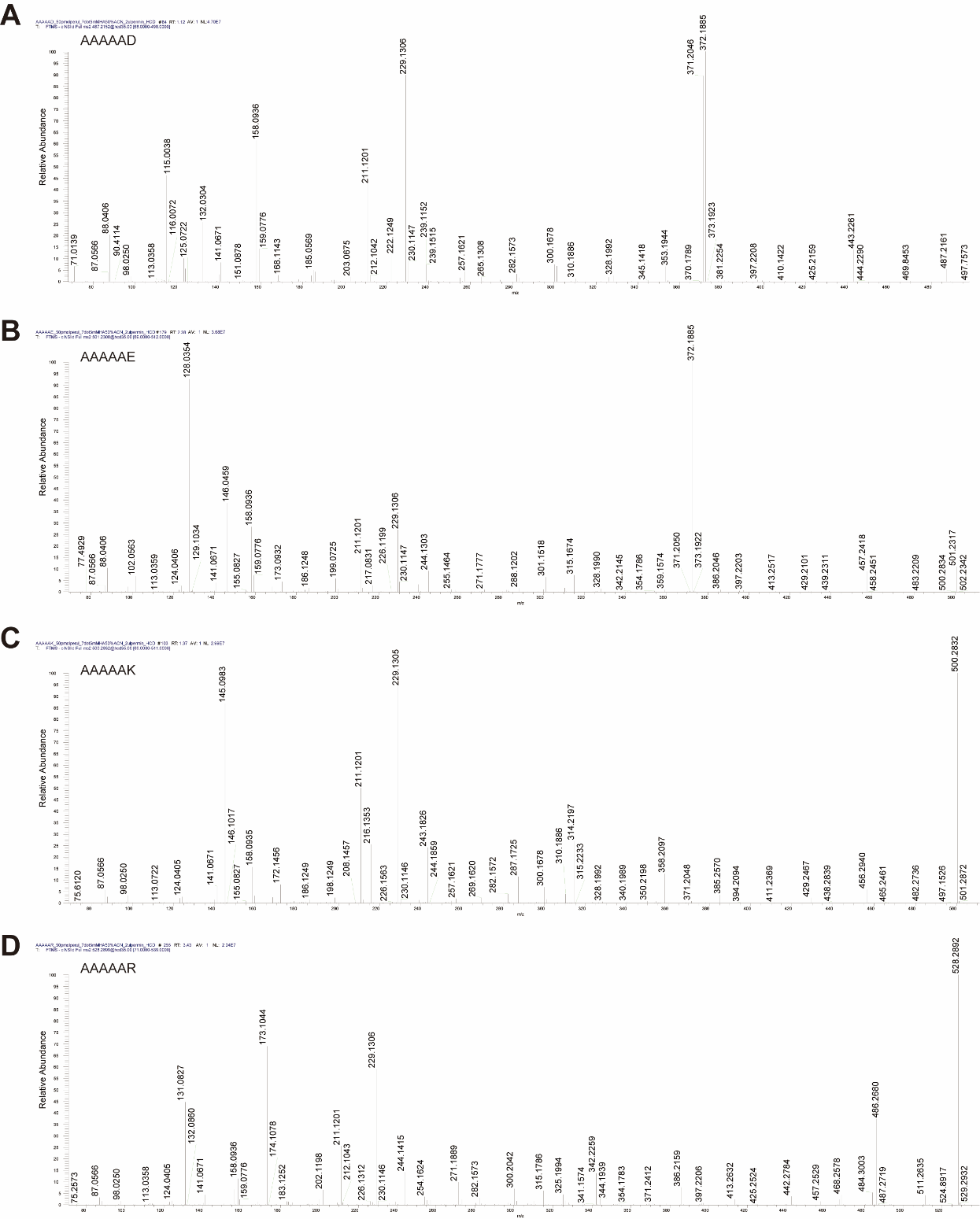

Fig. S1. Original (resolution of 120K) spectra of AAAAAD (A), AAAAAE (B), AAAAAK (C), and AAAAAR (D) at NCE 35.

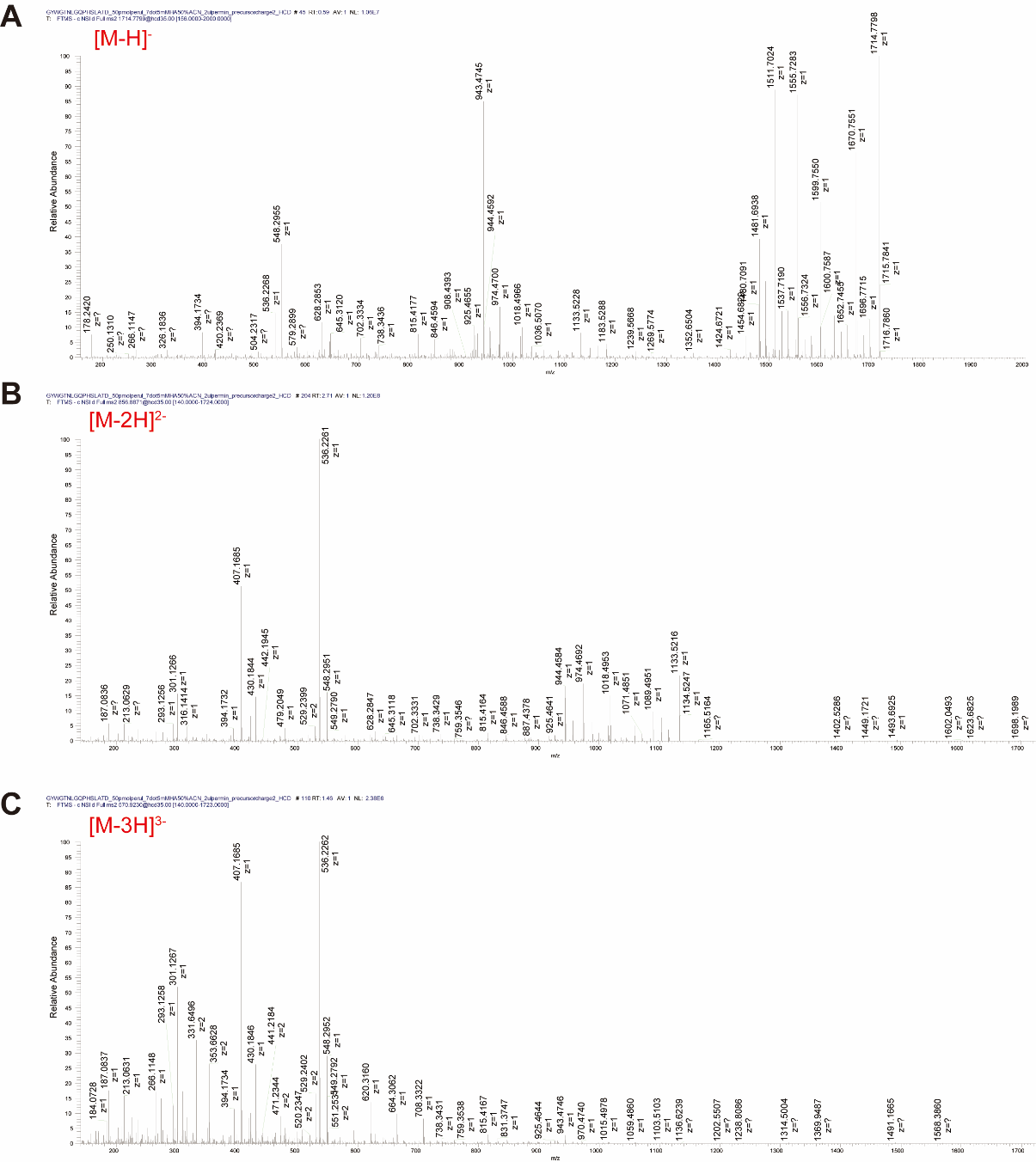

Fig. S2. Original HCD (resolution 120K) spectra of GYWGTNLGQPHSLATD from a 1^-^ (A), 2^-^ (B), or 3^-^ (C) precursor.

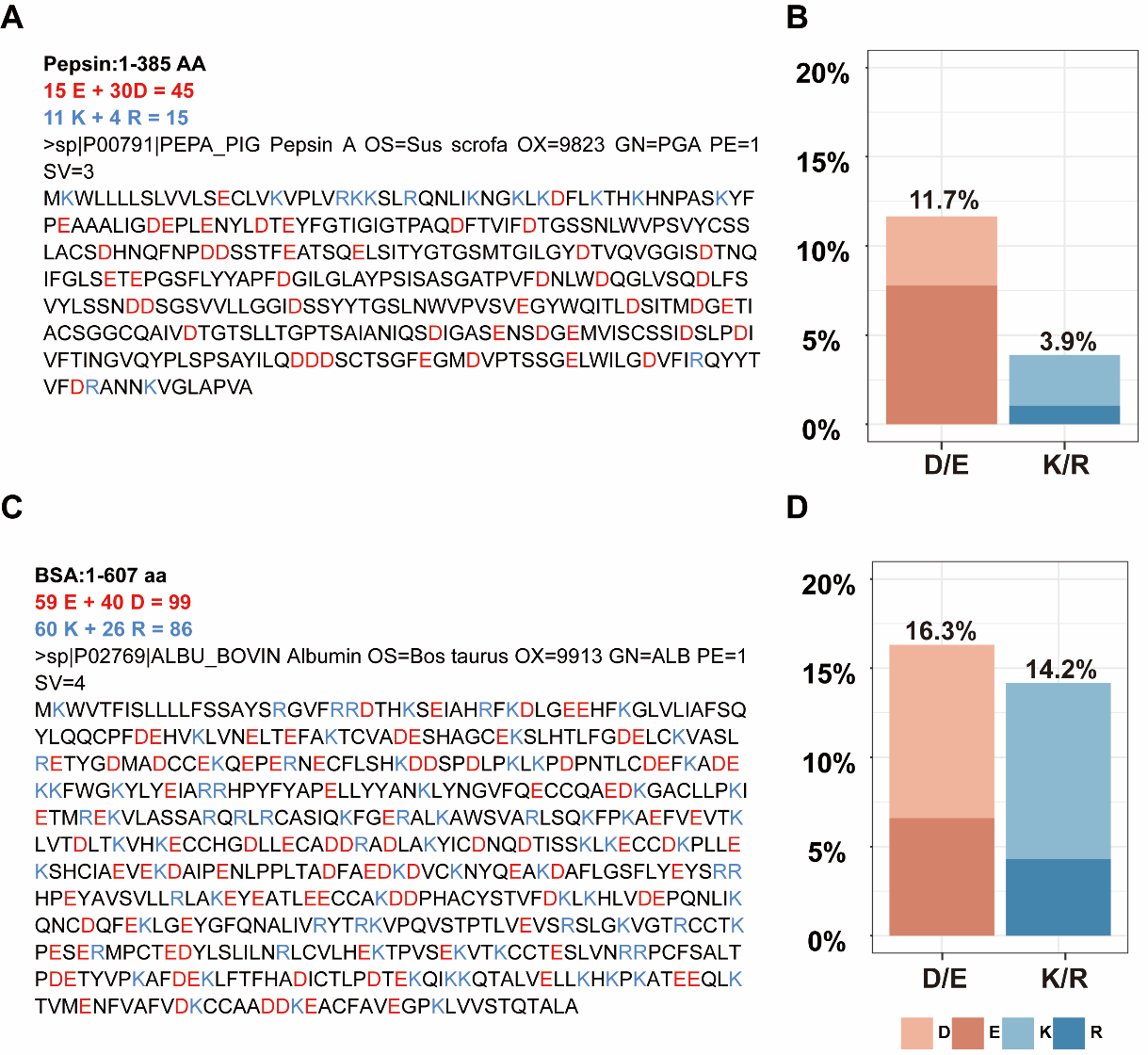

Fig. S3. The sequence (A, C), percentage of D+E and K+R (B, D) of pepsin (A, B) and BSA (C, D). The acidic amino acid are highlighted in red and the basic amino acids are highlighted in blue.
